# Mining downy mildew susceptibility genes: a diversity study in grapevine

**DOI:** 10.1101/2020.01.15.898700

**Authors:** Carlotta Pirrello, Tieme Zeilmaker, Luca Bianco, Lisa Giacomelli, Claudio Moser, Silvia Vezzulli

**Affiliations:** Research and Innovation Centre, Edmund Mach Foundation, Via E. Mach 1, 38010 San Michele all’Adige, Italy; Department of Agricultural, Food, Environmental and Animal Sciences, University of Udine, via delle Scienze 206, 33100 Udine, Italy; SciENZA Biotechnologies B.V., Sciencepark 904, 1098 XH Amsterdam, The Netherlands

**Keywords:** Amplicon sequencing, disease resistance, DLO, DMR, SNP, *Vitis* spp

## Abstract

Several pathogens continuously threaten viticulture worldwide. Until now, the investigation on resistance loci has been the main trend to understand the interaction between grapevine and mildew causal agents. Dominantly inherited gene-based resistance has shown to be race-specific in some cases, to confer partial immunity and to be potentially overcome within a few years since its introgression. Recently, on the footprint of research conducted on Arabidopsis, the putative hortologues of genes associated with downy mildew susceptibility in this species, have been discovered also in the grapevine genome. In this work, we deep-resequenced four putative susceptibility genes in 190 highly genetically diverse grapevine genotypes to discover new sources of broad-spectrum recessively inherited resistance. The scouted genes are *VvDMR6-1, VvDMR6-2, VvDLO1, VvDLO2* and predicted to be involved in susceptibility to downy mildew. From all identified mutations, 56% were Single Nucleotide Polymorphisms (SNPs) in heterozygosity, while the remaining 44% were homozygous. Regarding the identified mutations with putative impact on gene function, we observed ~4% genotypes mutated in *VvDMR6-1* and ~8% mutated in *VvDMR6-2*, only a handful of genotypes that were mutated in both genes. ~2% and ~7% genotypes showed mutations in *VvDLO1* and *VvDLO2* respectively, and again a few genotypes resulted mutated in both genes. In particular, 80% of impacting mutations were heterozygous while 20% were homozygous. The current results will inform grapevine genetics and corroborate genomic-assisted breeding programs for resistance to biotic stresses.

**Significance statement:** A survey on the genetic diversity of downy mildew susceptibility genes in grapevine varieties and wild species reveals a potential valuable for genomic-assisted breeding as well as tailored gene editing to induce disease resistance.

## INTRODUCTION

Crop plants encounter constant biotic challenges, and these threats have been commonly managed with pesticides and fungicides. Developing disease-resistant varieties is a convenient alternative to chemical control methods to protect crops from diseases. When a pathogen recognizes and invades the plant tissues and a plant-pathogen interaction is established, it faces the response of the host involving activation of signals that result in a rapid defence response. This immune response helps the host plant to avoid further infection of the disease (Gururani *et al.*, 2012). To suppress this immunity, pathogens produce effector molecules to alter host responses and support compatibility. In turn, plants evolved the ability to recognize these effectors by using resistance (R) genes. The majority of R-genes encode nucleotide-binding leucine-rich-repeat (NBS-LRR) proteins. Since R genes are specifically directed towards highly polymorphic effector molecules or their modifications, this kind of immunity is dominantly inherited, mostly race-specific and rapidly overcome by the capacity of the pathogen to mutate (Jones & Dangl, 2006). Analyses of whole-genome sequences have provided and will continue to provide new insights into the dynamics of R-gene evolution (Meyers, Kaushik, & Nandety, 2005).

Besides the established *R* gene model, the susceptibility (*S*) gene model has been more recently defined. All plant genes that facilitate infection and support compatibility can be considered *S* genes (reviewed in van Schie & Takken, 2014). They can be classified into the following three groups based on the point at which they act during infection: those involved in early pathogen establishment, those involved in modulation of host defences, and those involved in pathogen sustenance (Fawke, Doumane, & Schornack, 2015). The concept of susceptibility genes was first explored in barley by Jorgensen (1992) with the *MLO* (Mildew resistance Locus O) gene involved in susceptibility to powdery mildew. Later, *mlo* mutants were identified also in cucumber, melon, pea, tomato and tobacco (Kusch & Panstruga, 2017). Other analyzed susceptibility genes are the so called *DMR* (Downy Mildew Resistant) genes firstly characterized in Arabidopsis by Van Damme *et al.* (2005, 2008) and *DLO* (*DMR-*like Oxygenases) (K. Zhang, Halitschke, Yin, Liu, & Gan, 2013). Initially the Arabidopsis thaliana dmr6 mutant was isolated from an EMS population for its resistance to *Hyaloperonospora arabidopsidis*, the DM causal agent in this species (Van Damme *et al.*, 2005). Orthologs were readily identified in tomato (de Toledo Thomazella *et al.*, 2016) as well as many other crops (e.g. Schouten, Krauskopf, Visser, & Bai, 2014; Sun *et al.*, 2017) and fruit trees (e.g. Zeilmaker *et al.*, 2015; Zhang *et al.*, 2018). Mutations in DMR6 confer broad-spectrum resistance; *Sldmr6-1* tomato mutant plants show resistance against *Phytophthora capsici*; *Pseudomonas siringae* and *Xanthomonas* spp. (de Toledo Thomazella *et al.*, 2016).

In order to identify mutations and to deepen their impact on plant performance, studies of genetic diversity are essential and have been extensively performed in the plant kingdom, although compared to animals and humans their sequel is still in its infancy. A SNP (Single Nucleotide Polymorphism) provides the ultimate form of molecular marker, based on differences of individual nucleotide bases between DNA sequences (Ganal, Altmann, & Röder, 2009). SNPs are more abundant in the genome and more stably inherited than other genetic markers (Brookes, 1999) and they can be classified into random, gene targeted, or functional markers according to their localization (Andersen & Lübberstedt, 2003). The discovery of functional SNPs - that cause phenotype variations - is challenging and have been scarcely described in literature. In particular, functional SNPs were used to target flowering time and seed size in lentil (Polanco *et al.*, 2019), midrib colour in sorghum (Burow *et al.*, 2019), leaf hair number in turnip (Zhang *et al.*, 2018), grain length (Fan *et al.*, 2009) and blast resistance in rice (Yang *et al.*, 2017).

A variety of approaches have been adopted to identify novel SNPs (Edwards *et al.*, 2007). In the last decade, computational approaches have dominated SNP discovery methods due to the advent of Next Generation Sequencing (NGS, Varshney *et al.*, 2009), followed by the third-generation sequencing platforms (TGS, Schadt, Turner and Kasarskis, 2010), and the consequent ever-increasing sequence information in public databases. Since the first whole plant genome sequenced (The Arabidopsis Genome Initiative, 2000), *de novo* and reference-based SNP discovery and application are now feasible for numerous plant species. Large scale SNP discovery was performed in almost all sequenced plant genomes such as maize (Ching *et al.*, 2002), Arabidopsis (Atwell *et al.*, 2010), rice (Xu *et al.*, 2012), rapeseed (Raman *et al.*, 2014), potato (Vos *et al.*, 2015), and pepper (Hulse-Kemp *et al.*, 2016). On the method side, Genotyping-By-Sequencing (GBS) has recently emerged as a promising genomic approach to explore plant genetic diversity on a genome-wide scale (Peterson *et al.*, 2014), followed by the more cost-effective Genotyping-in-Thousands by sequencing (GT-seq) (Campbell, Harmon, & Narum, 2015). Genetic applications such as linkage mapping, phylogenetics, population structure, association studies, map-based cloning, marker-assisted plant breeding, and functional genomics continue to be enabled by access to large collections of SNPs (Kumar, Banks, & Cloutier, 2012). In parallel to SNP discovery based on whole genome sequencing, amplicon sequencing has also been successfully applied in plants (e.g. Durstewitz *et al.*, 2010; Yang *et al.*, 2016; Cho, Jones and Vodkin, 2017; Shimray *et al.*, 2017) although less frequently than in bacteria (e.g. Hong *et al.*, 2015) or viruses (e.g. Kinoti *et al.*, 2017).

Recently, as advocated by Gupta *et al.* (2001), progress has also been made in the development and use of SNPs in woody plants, including some crop and tree species as apple (Bianco *et al.*, 2016), walnut (Marrano *et al.*, 2019), sweet cherry (Hardner *et al.*, 2019), pear (X. Li *et al.*, 2019), and coffee (Merot-L’anthoene *et al.*, 2019). This phenomenon is due to the boost in the sequencing of cultivated plant genomes to provide high-density molecular markers for breeding programs aimed to crop improvement as well as to clear up evolutionary mechanisms through comparative genomics (Feuillet *et al.*, 2011; Bolger *et al.*, 2014). In grapevine a great deal of progress has been made from the first SNP identification in the pre-genomic-era (Owens, 2003) to the sequencing of the whole genome of several *Vitis vinifera* cultivars (Jaillon, 2007; Velasco *et al.*, 2007; Carrier *et al.*, 2012; Gambino *et al.*, 2017; Roach *et al.*, 2018) and to the very recent report of the genome sequence of *Vitis riparia* (Girollet, Rubio, & Bert, 2019). The latter represents a turning point on the scavenging of genomes that are donors of disease resistance. This issue in *Vitis* spp. is faced by identifying *R* loci, underlying *R* genes, through Quantitative Trait Loci (QTL) analysis in different genetic backgrounds. Nowadays, 13 *R* loci against powdery mildew and 27 to downy mildew have been identified with different origins; mainly from American and Asian wild species (Topfer and Hausmann, 2010).

Nowadays, a promising approach to cope with disease resistance is represented by the study of *S* loci. Based on a high-resolution map, Barba *et al*., (2014) identified on chromosome 9 a locus (*Sen1*) for powdery mildew susceptibility from ‘Chardonnay’, finding evidence for quantitative variation. Moreover, on the footprint of research conducted on model plants, genes associated with mildew susceptibility have been discovered and dissected also in the grapevine genome. 7 *VvMLO* orthologs in tomato and Arabidopsis were identified and members of *VvMLO* gene family showed transcriptional induction upon fungal inoculation (Winterhagen *et al.*, 2008; Feechan, Jermakow and Dry, 2009). Lately, a significant response in terms of powdery mildew resistance has been achieved by silencing of *VvMLO7* and *VvMLO6* through RNAi in grapevine (Pessina *et al.*, 2016).

In this research we aim to investigate the diversity of the DMR6 and DLO genes in a wide set of *Vitis* spp. to broaden our knowledge of the genetic variation present. This information will enhance our knowledge of possible alternative or integrative solutions compared to the use of *R* loci.

## RESULTS AND DISCUSSION

### Sequencing and mapping

In order to identify potentially disrupting mutations, coding sequences of the four *VvDMR6.1*, *VvDMR6.2, VvDLO1* and *VvDLO2* genes (**Table 1**) from 190 genotypes (**Table S1**) were deep-sequenced and mapped on the reference genome PN40024 12X V2 (see Materials and Methods section). Total sequence coverage of all genes together was 12,476,502 reads. *VvDMR6.1* was covered by 5,450,614 reads (44%), *VvDMR6.2* by 3,476,587 (28%)*, VvDLO1* by 3,270,318 (26%), and *VvDLO2* by 278,983 (2%). The highest coverage was detected in hybrid genotypes with a total of 9,357,649 reads (75%), followed by *vinifera* with 1,333,887 (11%), hybrids/wild species with 964,847 (8%) and wild species with 814,225 (6%).

**Table 1.**
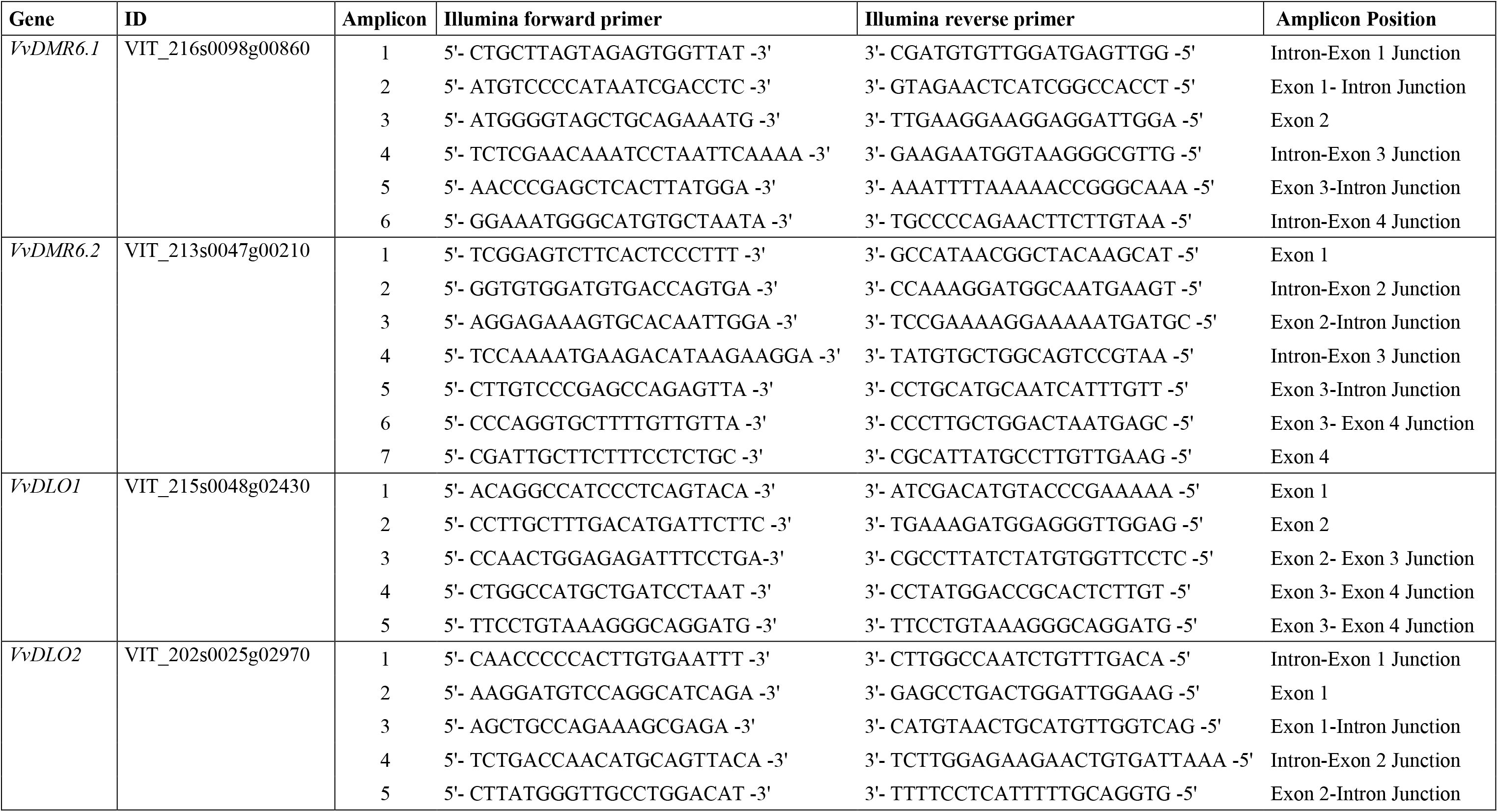
Targeted genes, amplicons with their genome positions and primers.

A total of 738 mutations were detected; 17 (~2%) short In/Dels and 721 point mutations, including heterozygous (56%) and homozygous (44%) SNPs.

### Genetic diversity assessment

Amplicons were classified according to their rate of polymorphism: from the most polymorphic VvDLO2_1 (~13% of the total mutations); to the ones carrying ~8% of mutations VvDMR6.1_3, VvDMR6.1_2, VvDMR6.2_3 gradually decreasing to the lowest rate of polymorphism (less than 3%) in VvDMR6.2_7 and VvDLO1_4.

Moreover, out of a total 738 mutations, 25 (~3.4%) triallelic variants were detected of which 13 in hybrids, 8 in wild species, 9 in *vinifera* varieties and 8 in hybrid/wild species. Triallelic mutations were mainly found in *VvDLO2* (12; ~1.6%) followed by *VvDMR6.1* (7; ~1%), *VvDMR6.*2 3 (~0.4%) and *VvDLO1*. As reported by Bianco *et al.* (2016) and Marrano *et al.* (2019), triallelic variants are usually discarded in SNP-based analyses to avoid incorrect genotypic information. Nevertheless, other authors provide data on their abundancy. The occurrence of the identified triallelism for each gene is consistent with previous work in grapevine (Lijavetzky *et al.*, 2007; Vezzulli *et al.*, 2008a; 2008b). In contrast, such a high representation of triallelic mutations in our accessions is due to the great genetic variability considered.

Considering the 696 biallelic mutations in all genotypes, 75% were transitions (A↔G, C↔T) and 25% were transversions (A↔C, A↔T, C↔G, G↔T) with a transition/transversion ratio of 3. Both *vinifera* varieties and hybrids show the same assortment with 77% transitions and 23% transversions, quite far from the ratio (~1.6) observed in the same taxa by Vezzulli *et al.* (2008a). In wild species the percentages were 73% and 27% respectively, while 71% and 29% were the values observed in hybrid/wild species genotypes. The current results slightly diverge from the usual transitions/transversions ratio found in grapevine (~1.5 in Salmaso *et al.*, 2004; Lijavetzky *et al.*, 2007; Vezzulli *et al.*, 2008a; 2008b; ~2 in Marrano *et al.*, 2017) as well as in beetroot (Schneider *et al.*, 2001), potato (Simko, Haynes, & Jones, 2006) and cotton (Byers *et al.*, 2012), while they are much higher than in soybean (Zhu *et al.*, 2003) and almond (Wu *et al.*, 2008).

SNP frequency was calculated as average and per gene for every taxon. *Vinifera* varieties showed the lowest average frequency (1 variant every 68.25 bp) with high differences between the target genes: 1 every 30.36 bp in *VvDMR6.1*, 1 every 46.09 bp in *VvDMR6.2*, 1 every 56.32 bp in *VvDLO1* and 1 every 140.22 bp in *VvDLO2.* A comparable polymorphism rate (1 SNP every 69 bp in coding regions) was found in both cultivated (spp. *sativa*) and non-cultivated (spp. *sylvestris*) *vinifera* species by Lijavetzky *et al.* (2007). In contrast, Vezzulli *et al.* (2008a) estimated 1 SNP every 117 bp in cultivated v*inifera* and 1 every 169 bp in wild *vinifera* individuals coding sequence. Moreover, in this study the detected average frequency was 1 variant every ~55 bp in both wild species and hybrid/wild species genotypes, while for the single genes they showed respectively 1 every 43.17 bp and 1 every 25.43 bp in *VvDMR6.1*, 1 every 50.70 bp and 56.33 bp in *VvDMR6.2*, 1 every 77.63 bp and 94.09 bp in *VvDLO1* and 1 every 45.52 bp and 49.86 bp in *VvDLO2.* Hybrids showed a higher average frequency (1 every 36.44 bp) due to the dramatically high frequency values in *VvDMR6.1* (1 every 13.41 bp) and in *VvDMR6.*2 (1 every 19.95 bp), 1 every 26.46 bp in *VvDLO1* and 1 every 85.92 bp in *VvDLO2.* Studying different *Vitis* spp. genotypes, Salmaso *et al.* (2004) observed an average of 1 SNP every 47 bp in the coding sequence of a set of genes encoding proteins related to sugar metabolism, cell signalling, anthocyanin metabolism and defence. Based on the first *Pinot noir* consensus genome sequence, the average SNP frequency was estimated at 4 SNPs every Kb (Velasco *et al.*, 2007), compatible with the use of such molecular markers for the construction of genetic maps in grapevine (Salmaso *et al.*, 2008). Higher polymorphism rates were found in other highly heterozygous tree species as peach (1 every 598 bp; Aranzana *et al.*, 2012), black cottonwood (1 every 384bp; Tuskan *et al.*, 2006), almond (1 every 114 bp; Wu *et al.*, 2008) and Tasmanian blue gum tree (1 every 45 bp; Thavamanikumar *et al.*, 2011).

As explained by Jones *et al.* (2007) and Grattapaglia *et al.* (2011), genotyping studies take advantage of different molecular markers, mostly relying on their informativeness. In this framework, SNPs are highly informative markers and this peculiarity is calculated as Minor Allele Frequency (MAF). SNPs are considered interesting for many goals when MAF values are >0.05 (Biswas *et al.*, 2015; Cheng *et al.*, 2019) but their main usefulness is due to the transferability across genotypes (>0.1; Lijavetzky *et al.*, 2007). In the current study, MAF was calculated for each biallelic mutation. MAF values 0.01≤x≤0.05 are represented by the 29% of mutations detected in total genotypes, in particular by the 23%, 0%, 2% and 3% in hybrids, wild species, *vinifera* varieties and hybrids/wild species, respectively. Values 0.05<x ≤ 0.1 are represented by 3% of the mutations in total genotypes and in wild species and by 2% in hybrids, *vinifera* varieties and hybrid/wild species. 0.1<x≤0.3 MAF values are represented by the 5% of mutations in total genotypes as in hybrids; wild species and *vinifera* varieties represented them by the 4% of their mutations and hybrid/wild species by the 2%. A very low percentage of mutations showed MAF 0.3<x≤0.5: 3% for total genotypes, hybrids and vinifera; 2% for wild species and hybrid/wild species. Finally, MAF >0.5 was very poorly represented by mutations in total genotypes and each taxon. SNP informativeness depends on their reliability among individuals and species and their high transferability rates probably are not consistent with a direct impact on the genetic sequence (when in coding regions). Considering previous studies in grapevine, a larger representativeness of MAF values <0.1 was found in non-*vinifera* genotypes and rootstocks, non-cultivated *vinifera* showed a MAF 0.05<x<0.3 while MAF >0.1 were severely represented by *vinifera sativa* (Lijavetzky *et al.*, 2007; Vezzulli *et al.*, 2008a; Emanuelli *et al.*, 2013; Marrano *et al.*, 2017). In the current study, the aim to focus on impacting mutations was achieved, since MAF ≤0.05 is a distinguishing mark for rare SNPs, which may be not considered interesting for SNP-arrays but which are most likely affecting the gene sequence and putatively protein activity.

### Mutation impact evaluation

In crops like tomato (Aflitos *et al.*, 2014) and *cucurbita* spp. (Xanthopoulou *et al.*, 2019), coding regions and whole genome sequence were scouted to find impacting mutations using SnpEff (Cingolani *et al.*, 2012). A non-synonymous/synonymous mutation ratio of ~1.5 was found in tomato cultivated cv. In *cucurbita* spp., the ratio was ~0.8 but only 9% of genetic variants showed HIGH or MODERATE impact in full genomic sequence, suggesting a great presence of intergenic mutations. In walnut tree genomic sequence, Marrano *et al.* (2019) identified 2.8% potentially impacting variants, while in the pear genome 55% of mutations were classified as missense and 1% with HIGH impact (Dong *et al.*, 2019).

In the current study, upon the variant discrimination performed according to their impact on codon sequence, 27% of total mutations (in particular, 27% in *VvDMR6.1*, 25% in *VvDMR6.2*, 30% in *VvDLO1* and 25% in *VvDLO2)* were classified as “MODIFIER”: falling into intronic regions or upstream/downstream the gene. “LOW” impact variants represented the 32% (36% in *VvDMR6.1*, 32% in *VvDMR6.2*, 32% in *VvDLO1* and 28% in *VvDLO2*), responsible for synonymous mutations or falling into splice regions. Of total mutations, 38% (in particular, 35% in *VvDMR6.1*, 40% in *VvDMR6.2*, 35% in *VvDLO1* and 43% in *VvDLO2*) brought to non-synonymous variants and were then classified with “MODERATE” impact. Percentages partially corroborated in *vinifera* by Amrine *et al.* (2015), with ~90% of MODIFIER and LOW mutations and ~8% non-synonymous variants in gene sequence. The lowest number of variants (3%: 2% in *VvDMR6.1*, 2% in *VvDMR6.2*, 3% in *VvDLO1* and 4% in *VvDLO2*) was classified with “HIGH” impact as being responsible for sequence frameshift or premature stop codon occurrence. A significantly lower presence (0.7%) of HIGH impacting variants was observed in Thompson Seedless cv. by Cardone *et al.* (2016). The current aim to detect potentially disrupting mutations finds support in the great frequency of HIGH- and MODERATE-impact variants compared to the aforementioned works on grapevine.

Following the filtering of mutations classified as “MODERATE” and “HIGH” (41%) in order to discriminate amino acid variants according to their conservation, these variants were further checked and mutants carrying different chemical/physical properties from the reference were chosen (see Materials and Methods section). Finally, results from both analyses on amino acid sequence were cross-referenced and a total of 19 mutations was elected as potentially affecting the protein structure: 5 in VvDMR6.1, 4 in VvDMR6.2, 4 in VvDLO1 and 6 in VvDLO2 (**Table S2**).

Given the predicted complementarity of AtDMR6 and AtDLO in salicylic acid catabolism (K. Zhang *et al.*, 2013; Y. J. Zhang *et al.*, 2017), particular interest in these results is given by the occurrence of impacting elected mutations in each one of the four scouted genes. This may allow the use of *VvDMR6* and *VvDLO* genes in different combinations to enhance the impact of such homozygous mutations and likely avoid complementary effects.

### Mutated DMR and DLO gene combinations

Of the studied genotypes, 55 showed at least one of the elected mutations: 37 hybrids, 2 *vinifera* varieties, 6 wild species and 10 hybrid/wild species. 73% of 55 genotypes showed mutations only in one gene: 13% in *VvDMR6.1*, 29% in *VvDMR6.2*, 7% in *VvDLO1* and 24% in *VvDLO2*, while 27% were double mutants within 6 gene combinations (**Table 2**). Frequencies of occurring mutation arrangement (consensus sequence) were calculated for each gene. Regarding *VvDMR6.1* one main mutations set was shared by 13% of genotypes (belonging to hybrid taxon). 46% and 19% of genotypes (both clusters with only hybrid individuals) showed two shared assortments for *VvDMR6.2.* Only one set in *VvDLO1* was shared by 15% of genotypes (all wild species) while three different *VvDLO2* sets were shared respectively by 13% (all hybrids), 13% (belonging to hybrid and wild species taxon) and 9% (hybrid and hybrid/wild species individuals) of genotypes. All other genotypes showed unique assortment of mutations.

**Table 2.**
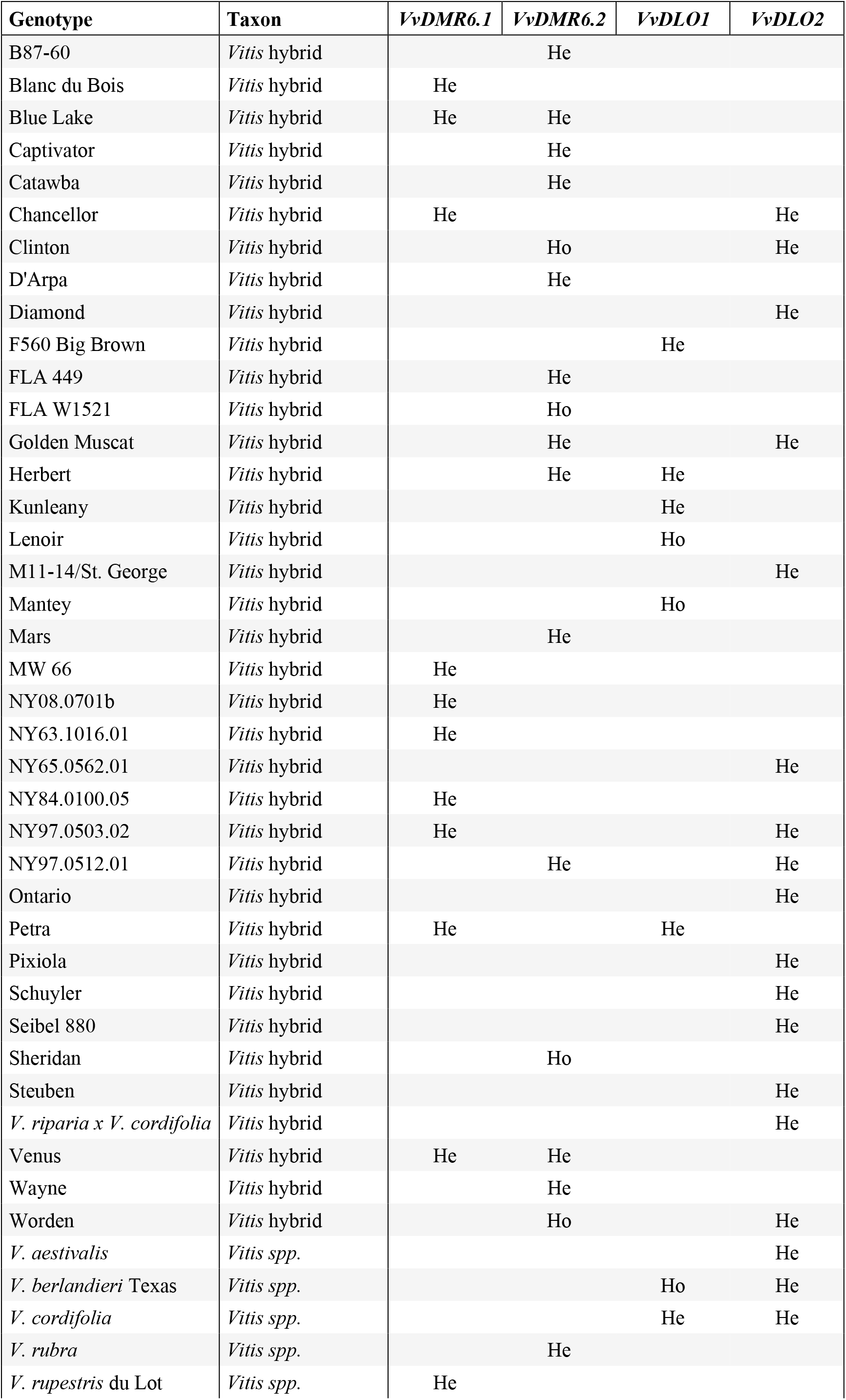
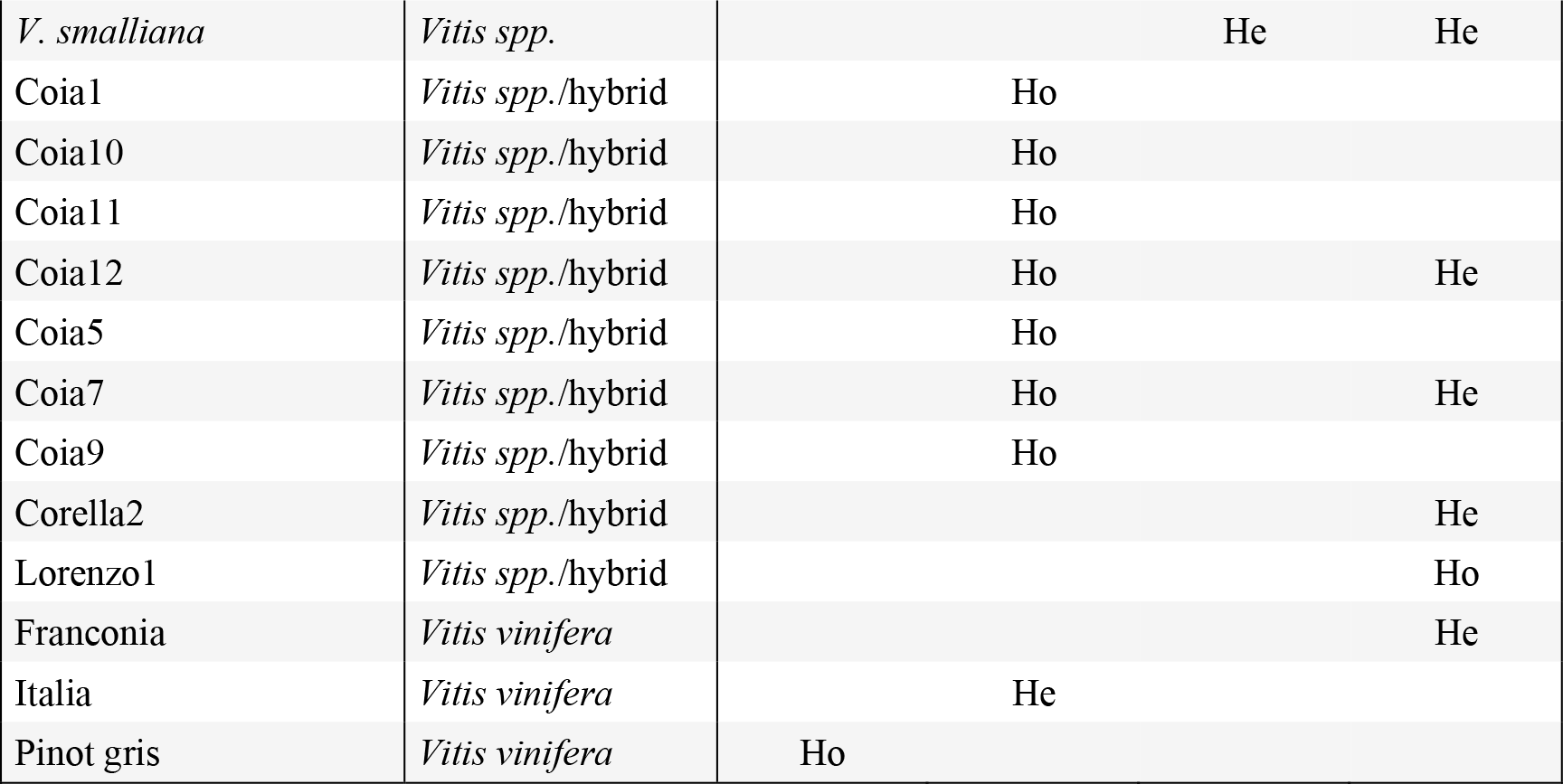
List of accessions carrying impacting mutations.

Induction of plant defence signalling involves the recognition of specific pathogen effectors by the products of specialized host *R* genes. Numerous plant *R* genes have already been identified and characterized and they are being efficiently used in crop improvement research programs (Gururani *et al.*, 2012). However, especially in tree species, selection of desirable resistant mutants come with a cost of lengthy and laborious breeding programs. The effort required to produce resistant plants is often baffled within a few years from the selection because the pathogen evolves mechanisms to circumvent the R-gene mediated immunity (Schaart *et al.*, 2016; Bisht *et al.*, 2019).

Exploitation of inactive alleles of susceptibility genes seems to be a promising path to introduce effective and durable disease resistance. Since S genes first discovery (Jorgensen, 1992), converting susceptibility genes in resistance factors has become the increasingly complementary strategy to that of breeding for R loci (van Schie & Takken, 2014), and the advent of new reliable genome editing tools has enhanced this trend. The use of genome editing technologies such as CRISPR-Cas9 allow to specifically and rapidly target susceptibility genes to indirectly obtain resistance in a chosen genetic background, which is highly desired in crops like grapevine where the genetic identity is economically important. However, generation of edited plants and testing of their phenotype still requires years (ffrench-Constant & Bass, 2017; Zaidi *et al.*, 2018). S genes may play different functions in the plant, thus pleiotropic effects associated with their knock-out may entail a certain fitness cost for the plant. Recently, quantitative regulation of gene expression has been achieved with genome editing on *cis-*regulatory elements (Rodríguez-Leal *et al.*, 2017; Wolter & Puchta, 2018; Bisht *et al.*, 2019) and this might be a strategy to limit negative drawbacks associated with a reduced S-gene function.

In this framework, thorough genetic diversity studies, as the one presented here, hold the potential to become a resource in different plant science contexts. The detection of specific homozygous variants in the natural pool can guide genome editing projects in targeting the “naturally” occurring mutations. This “tailored gene editing” mimicking natural polymorphisms, has been recently demonstrated by Bastet *et al.* (2017; 2019). Moreover, breeding programs could take advantage of the information on homozygous and heterozygous selected mutations of S-genes in a next-generation marker-assisted breeding program.

## MATERIALS AND METHODS

### Genetic material and target genes

In the current study, the four *VvDMR6.1*, *VvDMR6.2, VvDLO1* and *VvDLO2* genes were scouted in 190 grapevine genotypes (**Table 1, Table S1**). Out of these, 139 (73%) are *Vitis* hybrids, 28 (15%) are *V. vinifera* varieties, 12 (6%) belong to wild *Vitis* species and additional 11 (6%) are ascribed as hybrids/wild species.

### Amplicon sequencing and read processing

Genomic DNA was extracted from young grapevine leaves using DNeasy Plant Mini Kit (QIAGEN, Hilden, Germany.) according to the manufacturer’s protocol, then used to produce amplicons for deep-sequencing. PCR on the templates was performed using Phusion High-Fidelity Polymerase (NEB, Ipswich, Massachusetts, USA) according to the manufacturer’s protocol. Primers were specifically designed to amplify 250 bp of the coding regions of target genes and barcoded followed by in-house sequencing using the Illumina MiSeq platform (**Table 1**). A total of 19 amplicons was sequenced including six amplicons for *VvDMR6.1*, seven amplicons for *VvDMR6.2*, four amplicons for *VvDLO1* and two amplicons for *VvDLO2.*

Obtained amplicons were then mapped on the PN40024 12X reference genome (Jaillon, 2007) considering the latest V2 gene prediction (Vitulo *et al.*, 2014; Canaguier *et al.*, 2017) through Burrows-Wheeler alignment (BWA; Heng Li & Durbin, 2010) with no filter on mapping quality.

### Data mining

Variant calling was performed by BCFtools (H. Li *et al.*, 2009) using the following settings: minimum mapping quality 20; minimum genotype quality 20; minimum base quality 20; maximum per sample depth of coverage 1,000; minimum depth of coverage per site 10; keep read pairs with unexpected insert sizes (for amplicon sequencing). Filtering of results was done with VCFtools (Danecek *et al.*, 2011) to exclude all genotypes with quality below 20 and include only genotypes with read depth ≥ 10.

SnpEff was used to further discriminate variants according to their impact (MODIFIER, HIGH, MODERATE or LOW) on gene sequence (Cingolani *et al.*, 2012). Elected-impacting variants were then subject to SIFT (Sorting Intolerant From Tolerant) (P. Kumar, Henikoff, & Ng, 2009) analysis to assess the tolerance of aminoacidic variants on the protein primary structure, based on the alignment with sequences in SWISS-PROT/TrEMBL database. Only not tolerated mutations were considered for a last impact evaluation based on variants chemical-physical properties according to Betts & Russell (2003) (**Figure 1**). Both SnpEff and SIFT algorithms were used with default parameters settings.

**Figure 1.**
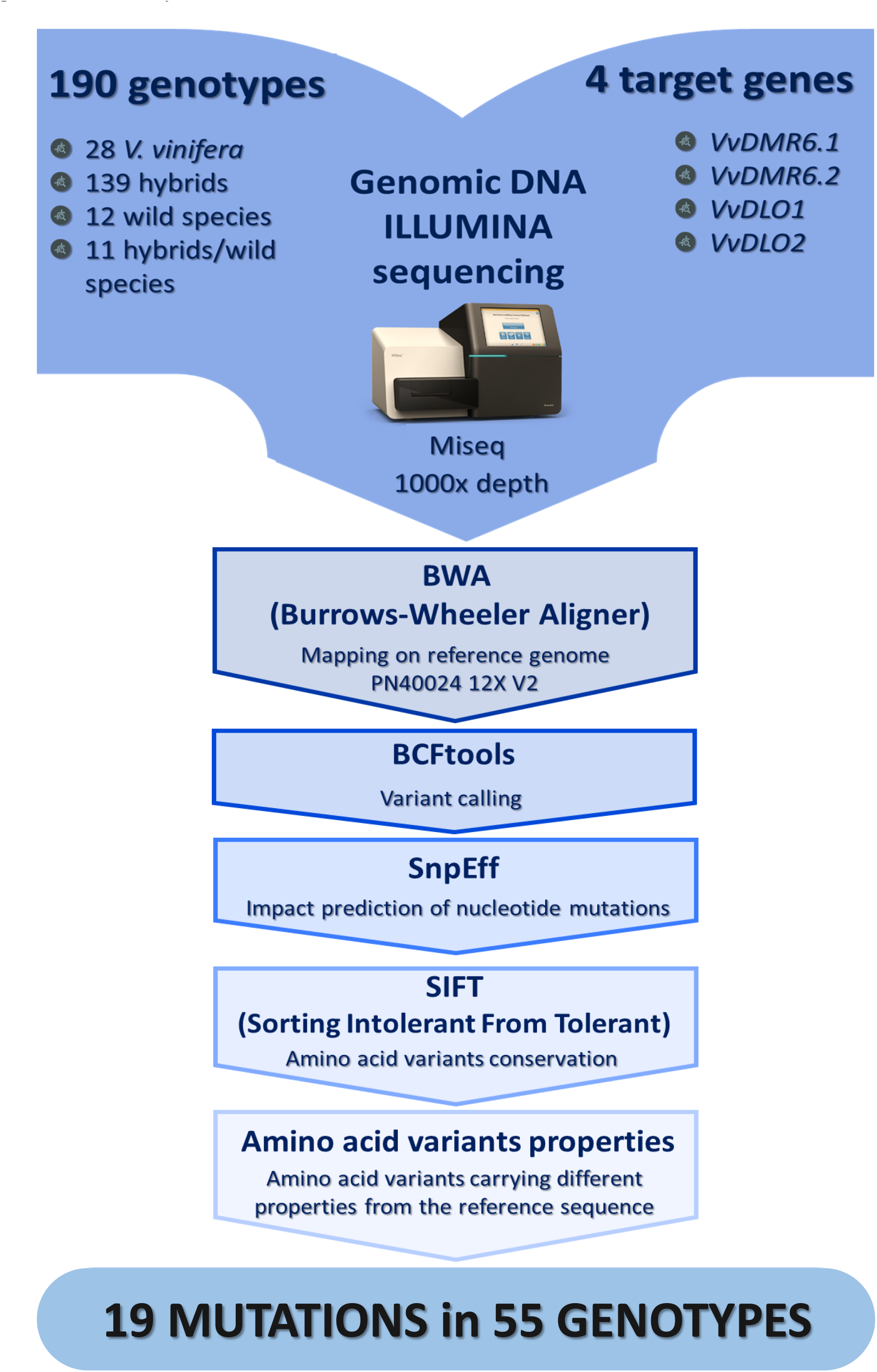
Data analysis flowchart.

### Statistical analysis

Data obtained from mapping and variant calling were dissected to extrapolate overall genetic information on the studied genotypes. Amplicons were classified according to their level of polymorphism. All the other parameters were calculated considering total accessions and the various taxon. For each gene, frequencies of occurring mutation arrangement were calculated along with mutation frequency, triallelic variants occurrence and MAF.

## Acknowledgments

CP acknowledges funding from a FEM PhD fellowship at University of Udine, including a support from SciENZA Biotechnologies bv.

